# TESOGENASE, An Engineered Nuclease Editor for Enhanced Targeted Genome Integration

**DOI:** 10.1101/2023.08.28.553855

**Authors:** Hangu Nam, Keqiang Xie, Ishita Majumdar, Shaobo Yang, Jakob Starzyk, Danna Lee, Richard Shan, Jiahe Li, Hao Wu

## Abstract

Non-viral DNA donor template has been widely used for targeted genomic integration by homologous recombination (HR). This process has become more efficient with RNA guided endonuclease editor system such as CRISPR/Cas9. Circular single stranded DNA (cssDNA) has been harnessed previously as a <u>g</u>enome engineering c<u>atalyst</u> (GATALYST) for efficient and safe targeted gene knock-in. However, the engineering efficiency is bottlenecked by the nucleoplasm trafficking and genomic tethering of cssDNA donor, especially for extra-large transgene integration. Here we developed enGager, <u>en</u>hanced <u>G</u>ATALYST <u>a</u>ssociated <u>g</u>enome <u>e</u>ditor system by fusion of nucleus localization signal (NLS) peptide tagged Cas9 with various single stranded DNA binding protein modules through a GFP reporter Knock-in screening. The enGager system assembles an integrative genome integration machinery by forming tripartite complex for engineered nuclease editors, sgRNA and ssDNA donors, thereby facilitate the nucleus trafficking of DNA donors and increase their active local concentration at the targeted genomic site. When applied for genome integration with cssDNA donor templates to diverse genomic loci in various cell types, these enGagers outperform unfused editors. The enhancement of integration efficiency ranges from 1.5- to more than 6-fold, with the effect being more prominent for > 4Kb transgene knock-in in primary cells. We further demonstrated that enGager mediated enhancement for genome integration is ssDNA, but less dsDNA dependent. Using one of the mini-enGagers, we demonstrated large chimeric antigen receptor (CAR) transgene integration in primary T cells with exceptional efficiency and anti-tumor function. These tripartite <u>e</u>ditors with <u>s</u>sDNA <u>o</u>ptimized <u>g</u>enome <u>en</u>gineering system (TESOGENASE^TM^) add a set of novel endonuclease editors into the gene-editing toolbox for potential cell and gene therapeutic development based on ssDNA mediated non-viral genome engineering.

**Highlight:**

- A reporter Knock-in screening establishes enGager system to identify TESOGENASE editor to improving ssDNA mediated genome integration
- Mini-TESOGENASEs developed by fusing Cas9 nuclease with novel ssDNA binding motifs
- mRNA mini-TESOGENASEs enhance targeted genome integration via various non-viral delivery approaches
- Efficient functional CAR-T cell engineering by mini-TESOGENASE

## INTRODUCTION

Clustered Regularly Interspaced Short Palindromic Repeats (CRISPR) and CRISPR-associated protein 9 (Cas9) is an effective and precise gene-editing tool that revolutionize diverse fields of biotechnology, medical and agricultural research (Cong et al., 2013; Doudna and Charpentier, 2014; Hsu et al., 2014). Cas9 protein combined with 20 nucleotides (nt) guide RNA finds and induces site specific double-stranded breaks (DSB) to the complementary region of the guide RNA for gene deletion, repair or insertion in prokaryotes and eukaryotes (Cong et al., 2013; Gilbert et al., 2013; Mali et al., 2013; Qi et al., 2013). Stemmed from this innovative tool, researchers around the world have developed many different applications of CRISPR/Cas9 such as gene knock-out using Non-Homogenous End Joining (NHEJ), correction of single point mutation by base editors that convert C to T or A to G nucleotide in a site specific manner, and targeted transcription regulation using catalytically dead Cas9 (dCas9) fused epigenetic regulators such as histone modifiers and DNA methylation modifiers (Cong et al., 2013; Hilton et al., 2015; Komor et al., 2016; Liu et al., 2016). However, targeted insertion of a new DNA sequence has been more challenging than the other CRISPR/Cas9 mediated applications.

Conventional targeted gene knock-in (KI) is achieved by homologous recombination (HR) which is extremely inefficient, especially for longer sequences. Homology directed repair (HDR), a highly regulated spatial and temporal cellular process induced by double stranded DNA break (DSB), significantly increases the targeted genome integration frequency. Currently, HDR is one of the most efficient ways to insert a few bases to 2Kb length of genes but is extremely inefficient for > 4Kb large payload (Mata López et al., 2020; Nishizono et al., 2020; Zhang et al., 2017).

To achieve efficient targeted genome integration, Adeno-Associated Virus Type 6 (AAV6) has been demonstrated as a prevalent donor carrier for HDR mediated genome engineering. For instance, AAV6 was used for efficient beta-globin gene integration in human and rodent hematopoietic progenitor cells as well as human stem cells (Dever et al., 2016; Martin et al., 2019; Wilkinson et al., 2021). However, recombinant AAV has been reported with safety issues and manufacturing challenges and succumbed for the use of large genome integration due to its packaging limit of ∼ 4.7 Kb (Dobrowsky et al., 2021; Fu et al., 2021; Nguyen et al., 2021). Non-viral targeted gene KI relies primarily on double-stranded DNA (dsDNA) and linear single-stranded DNA (lssDNA) donors. These nucleic acid donor templates have enabled relatively efficient gene targeting in various primary cells, including T cells and natural killer cells (Huang et al., 2021; Miura et al., 2015; Oh et al., 2022; Roth et al., 2018; Zhang et al., 2017). Recently, CD19 CAR-T cells engineered on an immune checkpoint PD-1 locus with linear dsDNA donor template have been reported to treat B cell Non-Hodgkin lymphoma (B-NHL) patients with desired safty and efficacy (Zhang et al., 2022). However, both forms of DNA template are not ideal HDR donor payloads. While dsDNA donors are easy to access, it succumbs from low editing efficiency and cGAS dependent cytotoxicity (Motwani et al., 2019; Roth et al., 2018). lssDNA has better performance for targeted gene integration but has major manufacture challenges (Hao et al., 2020). Most recently, circular single stranded DNA (cssDNA) has been demonstrated as a superior donor template for highly efficient genome targeting in mammalian cells (Iyer et al., 2022; Xie et al., 2022). However, non-viral DNA donor template mediated genome integration is largely bottlenecked by the nucleus delivery and genomic tethering of the donor DNAs, especially for targeted integration of large transgene payloads, such as polycistronic gene expression cassette. Therefore, developing a more efficient gene integration method, especially for long sequences with non-viral donor payload is an essential milestone to unlock the full potential for gene editing.

As HDR mediated genome integration is a highly regulated spatial and temporal process in nucleoplasm, the strategy to enhance the KI efficiency using DNA donor is 1) to efficiently deliver donor DNA into the nucleus and 2) to tether donor DNA onto the targeted genomic locus, both of which are critical steps to increase the active local concentration of the donor template to ensure effective homologous searching and pairing to complete the HDR process. This idea has been verified by multiple studies using different strategies. For instance, physical coupling of dsDNA donor templates together with Cas9 endonuclease by fusing Cas9 together with transcription factors or homologous recombinase such as Rad51, Rad52, POLD3 and the dominant negative form of 53BP1 has been shown to increase dsDNA mediated transgene KI (Gao et al., 2022; Jayavaradhan et al., 2019; Li et al., 2021; Lin-Shiao et al., 2022; Park et al., 2021; Rees et al., 2019; Reint et al., 2021; Shao et al., 2017, 2017; Tran et al., 2019). Other studies engineered the single or double stranded donor DNA with chemical modification such as biotinylation, 5’-triethylene glycol (TEG) modification or covalent conjugation of donor DNA with Cas9 endonuclease (Aird et al., 2018; Carlson-Stevermer et al., 2017; Ghanta et al., 2021; Ma et al., 2017; Savic et al., 2018). Similar strategy has been taken to create a sgRNA/donor DNA hybrid or a truncated Cas9 binding site on the donor DNA to achieve more efficient HDR mediated genome integration (Aiba et al., 2022; Nguyen et al., 2020; Shy et al., 2022; Simone et al., 2021). Overall, majority of the strategies focus on dsDNA as donor templates, with limited optimization on ssDNA donor templates.

We previously harnessed circular single stranded DNA (cssDNA) as a superior <u>g</u>enome engineering c<u>atalyst</u> (GATALYST) for efficient and safe targeted gene KI across multiple cell types to many targeted loci in the mammalian genome (Xie et al., 2022). Here we hypothesize that coupling GATALYST donor template with endonuclease editor as a nuclease/sgRNA/ssDNA tripartite complex can effectively increase the spatial and temporal enrichment of ssDNA donor template in the targeted genome microenvironment, thereby enhance the ssDNA mediated genome integration. Using a functional screening method based on a GFP reporter KI, we developed a set of <u>en</u>hanced <u>G</u>ATALYST <u>a</u>ssociated <u>g</u>enome <u>e</u>dito<u>r</u>s (enGager) by fusion of nucleus localization single (NLS) peptide tagged wild type Cas9 together with various single stranded DNA binding protein modules. These enGagers form tripartite complex with sgRNA and cssDNA donors as integrative genome integration machinery, thus facilitate the nucleus translocation and increase the active local concentration of donor DNA. When applied for targeted genome integration with cssDNA donor templates to diverse genomic loci in various cell types, these enGagers outperform unfused canonical endonuclease editors, in plasmid, mRNA and protein forms. The enhancement of integration efficiency ranges from 1.5- to more than 6-fold, with the effect being more prominent for larger than 4Kb transgene KI and in primary cells. These rationally designed tripartite <u>e</u>ditors with <u>s</u>sDNA <u>o</u>ptimized <u>g</u>enome <u>en</u>gineering system (Tesogenase^TM^) add a set of novel endonuclease into the gene-editing toolbox for potential cell and gene therapeutic development based on ssDNA mediated non-viral genome engineering.

## RESULTS

### Fusion of Cas9 and homologous recombination proteins enhance the ssDNA mediated knock-in

We and others have demonstrated that cssDNA acts as genome engineering catalyst (GATALYST) to enable safe and efficient targeted gene knock-in, but the knock-in efficiency for > 4Kb large transgene is relatively low (Iyer et al., 2022; Xie et al., 2022). To further optimize the cssDNA mediated gene knock-in, we sought to develop a set of enhanced GATALYST associated genome editors (enGager) that could form an endonuclease/sgRNA/ssDNA tripartite complex by physically recruiting the cssDNA donors, and facilitate the nucleus shuttling of the DNA donor by virtue of the nucleus localization signal (NLS) peptides tagged on the endonuclease editors. To develop such enGagers, we initially utilized an all-in-one (AIO) plasmid construct made by Cong et al to engineer dual NLS peptides tagged spCas9 fusion constructs for a GFP reporter screening (Cong et al., 2013). These enGager constructs contain bi-cistronic expression cassettes for Cas9-fusion endonuclease proteins and a single-guide RNA (sgRNA) targeting against the human *RAB11A* locus (Roth et al., 2018). We initially tested the Cas9 fusion with full length *E. coli* RecA and its human homolog protein Rad51 (Chen et al., 2008; Lin et al., 2006; Shinohara et al., 1992; Su et al., 2016). For potential better performance of Rad51-Cas9 fusion, we used AE and SEAD, two mutant forms of Rad51 with enhanced recombigenic activity identified previously (Rees et al., 2019). As the full length RecA and Rad51 have both recombinase activity and DNA binding capability, to uncouple these two functions, we also engineered an AIO construct by fusion of Cas9 with a 36 amino acid (AA) peptide “Brex” which is reported as a RecA binding motif to recruit endogenous RecA/Rad51 recombinase, but does not have DNA binding capability in the cells (Fig. 1A) (Ma et al., 2020). To directly compare the knock-in efficiency with various AIO enGager constructs, 2 ug of various size of GFP reporter cssDNA donor templates flanked with 306 nt 5’ homology arm and 315 nt 3’ homology arm targeting *RAB11A* locus were tested by co-electroporation delivery in K562 cells (Fig.1B). As illustrated in Fig. 1C, when DNA donor templates were co-electroporated with AIO constructs consisting of Cas9 fusion enGagers and *RAB11A* sgRNA, precise HDR of the locus can be evaluated by GFP reporter expression using flow cytometry. Day 3 post-electroporation, knock-in efficiency as measured by GFP+ cell percentage was much higher for Cas9-RecA, Cas9-Rad51 (AE) and Cas9-Rad51 (SEAD) enGager constructs (11.5%-15.4%) than Cas9 only construct (6.8%) for a 2 µg of cssDNA donor with 2Kb payload (Fig.1C). In contrast, no knock-in efficiency enhancement was observed with Cas9-Brex fusion construct. Interestingly, the KI efficiency by Cas9-Brex fusion nuclease is even lower than Cas9 alone (4.9% vs 6.8%), indicating that deploying the recombination function alone is not sufficient to potentiate cssDNA mediated genome targeting. We found that the enGager induced Knock-in enhancement is durable over time at 8 days and 14 days post-electroporation. In particular, the Cas9-RecA fusion enGager is able to induce >3-fold KI enhancement as compared to unfused Cas9 (Fig.1D). We further tested the knock-in efficiency at *RAB11A* locus in K562 cells for a larger cssDNA donor with a 4Kb GFP payload. EnGagers Rad51(AE), Rad51(SEAD) and RecA induced GFP KI enhancement by 1.78- to 2.35-fold compared to Cas9 alone (Fig.1E&F). Consistently, the Cas9-Brex fusion does not produce benefit for 4 Kb large payload KI.

**Fig.1.**
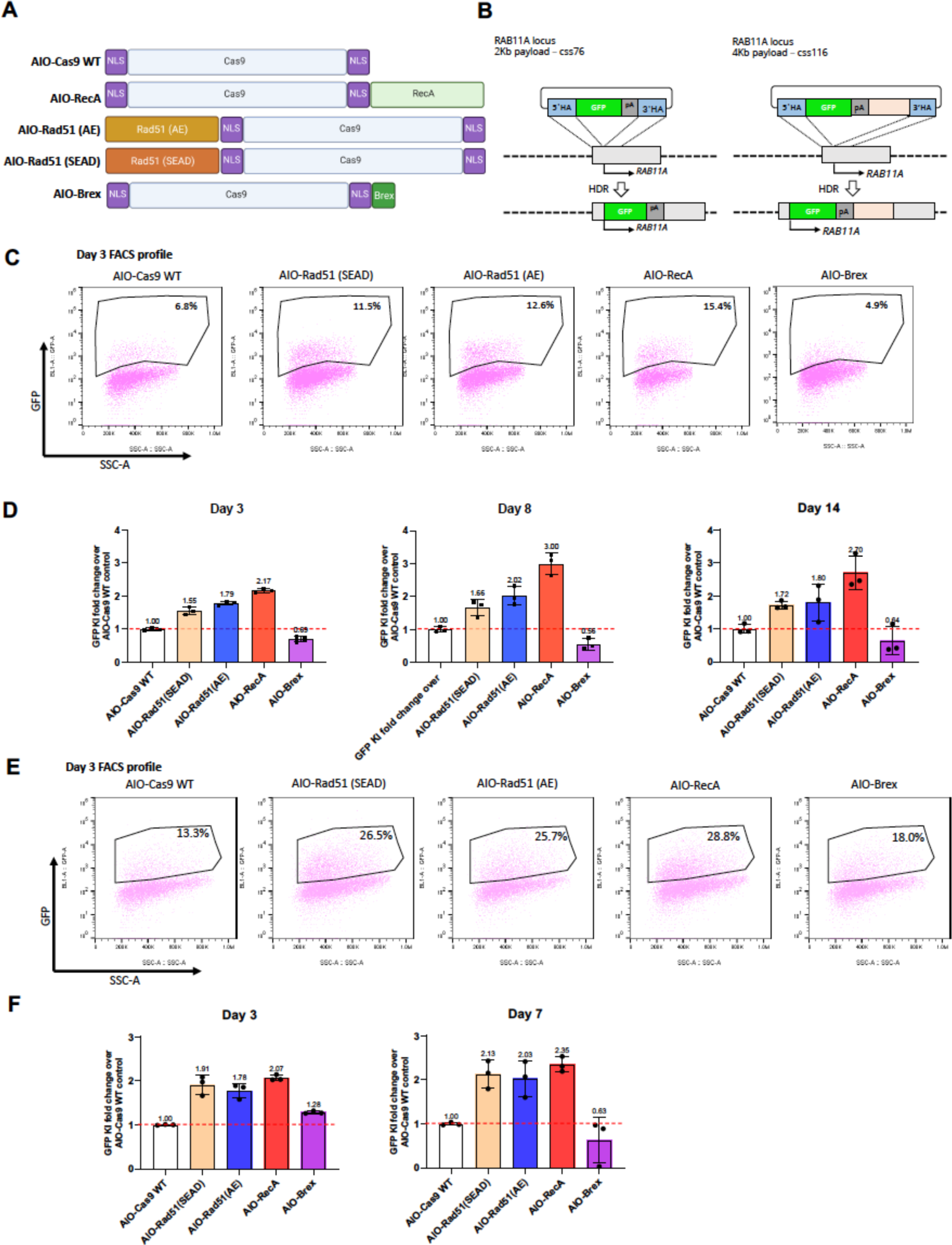
Fusion of Cas9 and homologous recombination proteins enhance the ssDNA mediated knock-in. A, Schematic diagram of various Cas9-homologous recombination protein fusion constructs (enGagers) in all-in-one (AIO) plasmid format modified from Addgene plasmid #42230. Two nuclear localization signals were added to the N’ and C’-termini of the Cas9 protein. RecA is the bacteria homologous DNA repair protein and Rad51(AE)/Rad51(SEAD) are two mutant variants from eukaryotes. Brex is a 36 amino acid peptide reported to recruit Rad51 in mammalian cells. B, Schematic diagram of Knock in strategy of a 2Kb (left) and 4Kb (right) cssDNA donor template for RAB11A locus. C, representative FACS profiles with gating strategy showing % of GFP transgene cassette Knock in on RAB11 locus at day 3 post electroporation for various enGagers listed in A. D, Quantification of 2Kb GFP transgene cassette Knock in fold change of various enGagers as compared to Cas9 WT at day 3 (left), 8 (middle) and 14 (right) post electroporation. E, representative FACS profiles with gating strategy showing % of 4Kb GFP transgene cassette Knock in on RAB11 locus at day 3 post electroporation for various enGagers listed in A. F, Quantification of 4Kb GFP transgene cassette Knock in fold change of various enGagers as compared to Cas9 WT at day 3 (left), 7 (right) post electroporation. Note that Brex enGager does not enhance knock in efficiency. Rad51 mutants and RecA enGagers increase both small and large transgene cassette knock in by 1.57 to 3.04-fold. RecA enGager outperforms among the enGagers tested. Bars represent mean ± SD from 3 biological replicates.

### Engineering mini-TESOGENASE with Cas9 fused ssDNA binding motifs

Based on the Brex- and full-length recombinase-Cas9 fusion results, we hypothesize that the rate-limiting step to enhance the ssDNA mediated HDR is to increase the effective nucleoplasm delivery of the donor template, which in turn increase the local concentration of the tripartite HDR machinery on the targeted genomic locus including the DNA donor template. Several seminal biochemical studies on RecA family DNA recombinase have identified an evolutionarily conserved ssDNA binding L2 loop peptide that encompasses 20-24 amino acid with an extremely conserved central aromatic residue (Maraboeuf et al., 1995, p. 2; Shinohara et al., 1992, p. 2; Sugimoto, 2000; Sugimoto and Nakano, 1997, p. 2; Voloshin et al., 1996, p. 2). To test our hypothesis, we engineered smaller enGager editors that only capture the DNA binding feature. We selected the 20 amino acid peptide sequences FECO, WECO and YECO from Voloshin et al which demonstrated several fold higher binding affinity to ssDNA than to dsDNA in sequence independent manner (Voloshin et al., 1996). We engineered smaller AIO constructs with Cas9 fused to either one copy or three tandem copies of these DNA binding motifs linked with multiple GS linker sequences (Fig.2A). When co-electroporated together with 2 ug of 2 Kb cssDNA *RAB11A* GFP donor template into K562 cells, these newly designed smaller enGagers demonstrated comparable level of enhanced knock-in efficiency to RecA full length enGager. Except for YECO3X construct, the fold changes range from 1.40 to 1.60, whereas the RecA enGager enhances 1.57-fold KI compared to Cas9 alone (Fig.2C&D). Interestingly, adding multiple copies of the peptide motif did not further enhance the KI efficiency for this 2 Kb cssDNA *RAB11A* GFP donor payload. These data validated the hypothesis that just installing the sequence independent DNA binding capability to nuclease editors could effectively stimulate the homology recombination mediated gene insertion. By doing so, we also discovered a set of smaller rationally designed enGagers that could be powerful editing tools for genome writing.

**Fig.2.**
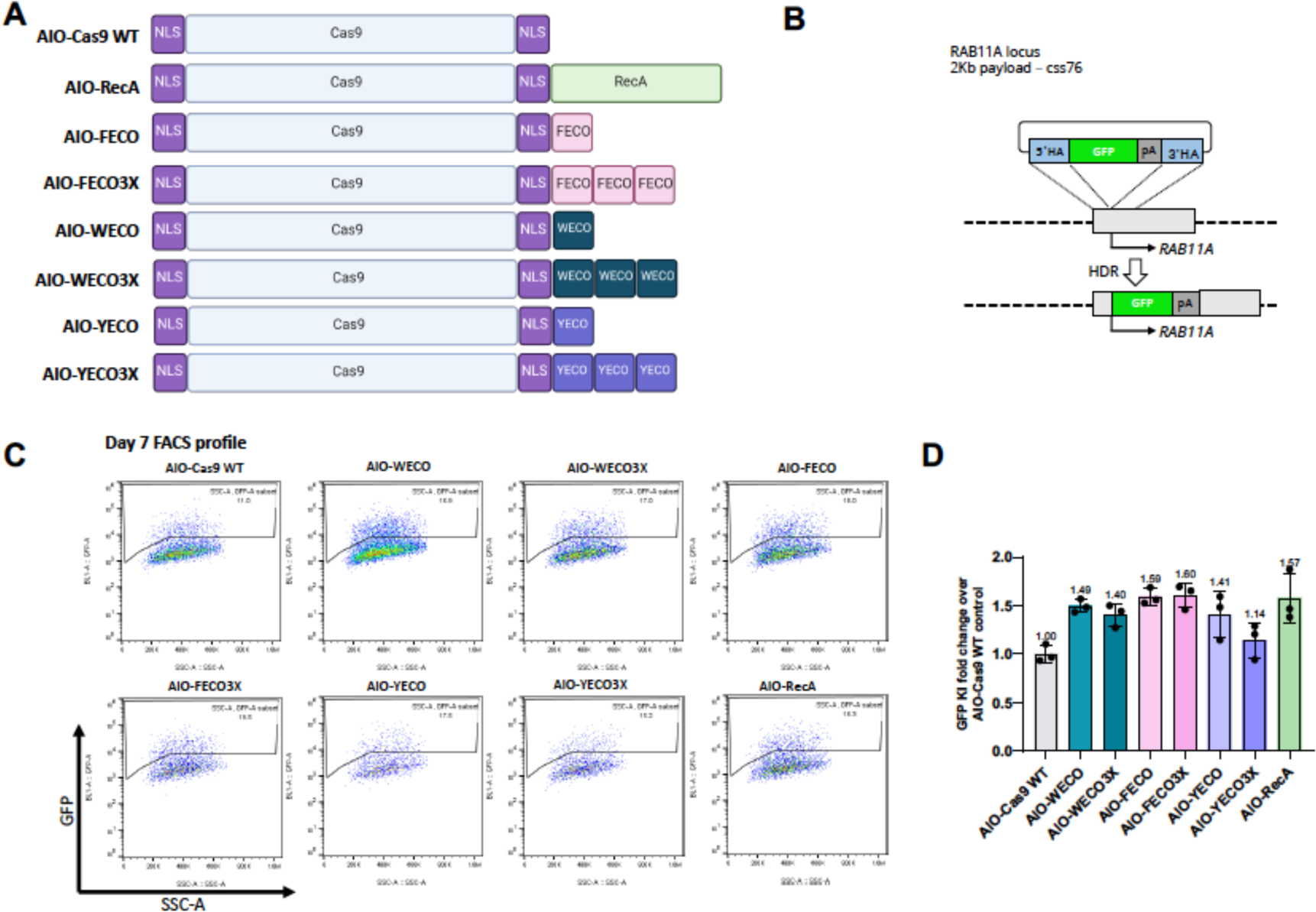
Identification of mini TESOGENASE with Cas9 fused ssDNA binding motifs in K562 cells. A, Schematic diagram of various Cas9-ssDNA binding motifs fusion constructs (enGagers) in all-in-one (AIO) plasmid format modified from Addgene plasmid #42230. Two nuclear localization signals were added to the N’ and C’-termini of the Cas9 protein. Cas9-RecA fusion construct was used as a positive control. FECO, WECO and YECO are 20 amino acid sequences previously identified as ssDNA binding motifs in various bacteria species of RecA(Voloshin et al., 1996), FECO3X, WECO3X and YECO3X are 3 tandem copies of the 20 aa peptides separated by multi-GS peptide linkers. B, Schematic diagram of Knock in strategy of a 2Kb cssDNA donor template for RAB11A locus. C, representative FACS profiles with gating strategy showing % of GFP transgene cassette Knock in on RAB11 locus at day 7 post electroporation for various small enGagers listed in A. D, Quantification of 2Kb GFP transgene cassette Knock in fold change of various enGagers as compared to Cas9 WT at day 7 post electroporation. Note that Cas9-FECO fusion performs similarly with Cas9-RecA fusion in cssDNA mediated transgene integration (1.59-vs 1.58-fold). EnGagers with 3X tandem ssDNA binding peptides do not further enhance knock in efficiency does not enhance knock in efficiency. Bars represent mean ± SD from 3 biological replicates.

### Identification of additional mini-TESOGENASE from Cas9-SSB module chimeras

To engineer a bigger collection of AIO enGager constructs We then did an extensive search of RecA family recombinase and other protein families that has ssDNA binding capability, either small motif peptide or structured protein domain. We included DrRecA full length and its 20 amino acid L2 peptide (Slade et al., 2009a; Zahradka et al., 2006), E.coli ssDNA binding protein (Meyer and Laine, 1990), Lambda Red subunit of bacteriophage recombineering protein (Mosberg et al., 2010), E.coli ssDNA annealing protein RecT (Noirot and Kolodner, 1998), archaea recombinase homolog RadA and RadB (Seitz et al., 1998; Wardell et al., 2017). (Fig.3A&B). These fusion constructs were screened using the 2 Kb cssDNA RAB11A GFP payload in K562 cells. Whereas Cas9-fusion with a ssDNA annealing protein RecT, the archaea homologs of EcRecA, RadA and RadB perform no better than Cas9 alone, all the other fusion constructs outperform Cas9 WT by 1.79- to 2.43-fold without compromising the cell viability (Fig.3C&D). Interestingly, the enGager with DrRecA full length fusion stimulates *RAB11A* GFP KI by 2.17-fold and the homologous 20 amino acid L2 peptide from DrRecA further improve the KI efficiency level to 2.43-fold of that of WT Cas9. As a comparison, FECO and RecA enGagers increase KI efficiency by 1.59- and 1.87-fold respectively. It is worth noting that *Deinococcus Radiodurans* Bacterium is one of the most radiation-resistant organism known to date due to its exceptional DNA repair capability by its DNA recombinase DrRecA (Slade et al., 2009b; Zahradka et al., 2006). We reasoned that the L2 loop peptide from the whole RecA phylogeny can be good candidate fusion peptides for small version of enGagers to stimulate HDR mediated genome integration. Fig.3E highlighted a few RecA L2 peptide sequences from archaea bacteria, E.coli and mammalian organism with a central amino acid with aromatic ring.

**Fig.3.**
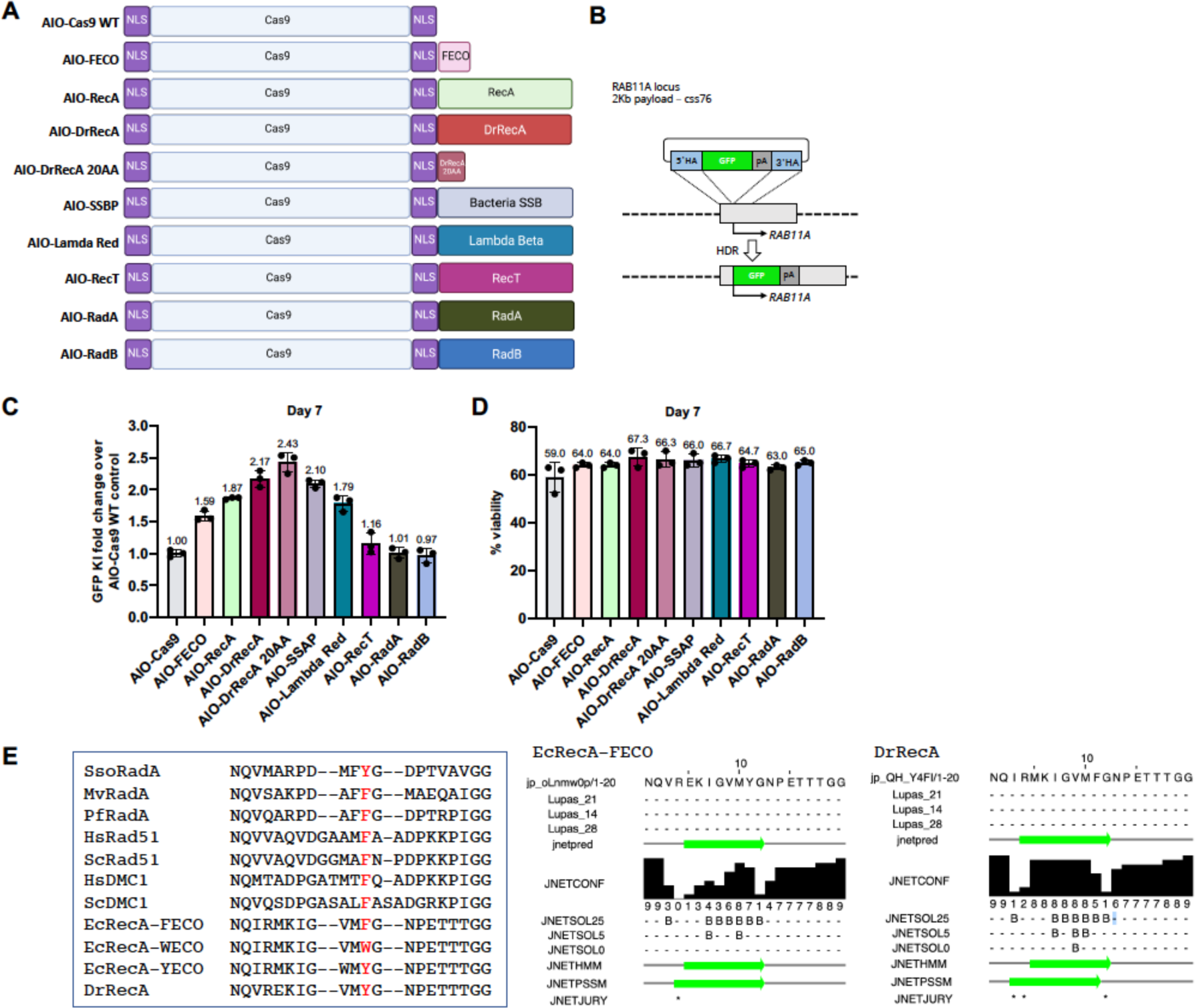
Identification of additional TESOGENASE from Cas9-ssDNA binding module chimeras in K562 cells. A, Schematic diagram of various Cas9-ssDNA binding protein and peptide fusion constructs (enGagers) in all-in-one (AIO) plasmid format modified from Addgene plasmid #42230. Two nuclear localization signals were added to the N’ and C’-termini of the Cas9 protein. Cas9-RecA and Cas9-FECO fusion constructs were used as positive controls. The fusion protein or peptides include DrRecA, 20 aa motif identified from DrRecA, SSAP, Lambda Red, RecT, RadA and RadB from Archaea. B, Schematic diagram of Knock in strategy of a 2Kb cssDNA donor template for RAB11A locus. C, Quantification of 2Kb GFP transgene cassette Knock in fold change of various enGagers as compared to Cas9 WT at day 7 post electroporation. Note that Cas9-DrRecA and Cas9-DrRecA20AA fusion has the highest performance in knock in with 2.17- and 2.43-fold as compared to Cas9 WT, respectively. D. Quantification of cell viability day 7 post electroporation. Bars represents mean ± SD from 3 biological replicates. E. Amino acid sequence alignment of 20AA of multiple E.coli RecA mutant variants and RecA from archaea and mammalian organism. Dr: *Deinococcus radiodurans*; Ec: *Escherichia coli; Sc: Saccharomyces cerevisiae; Hs: Homo sapiens; Pf: P. furiosus; Sso: S. solfatarcus*.

### Genome integration enhancement by TESOGENASE is ssDNA dependent

mRNA has been an attractive approach to reduce cytotoxicity and enhance delivery using lipid nanoparticle technology for vaccine development and gene therapy (Hou et al., 2021). Hereafter, except otherwise mentioned, we used mRNA form of RecA and FECO enGagers for the remaining of this study (Fig.4A). We initially validated the RecA enGager mediated enhancement of a 2Kb *RAB11A* GFP reporter integration in K562 cells by electroporation delivery. When co-electroporated with various dose of cssDNA donor templates at 0.3, 1, 1.5, 2 to 3 ug per reaction, RecA mRNA enGager (at 1 ug per reaction) significantly increases the GFP KI compared to WT Cas9 mRNA (Fig.4B&C). As RecA L2 peptides also have low binding affinity to dsDNA, we sought to investigate if these enGagers can facilitate HDR mediated KI by dsDNA donor templates. Using co-electroporation of 1 ug mRNA enGagers/sgRNA and 1 ug of 2Kb *RAB11A* GFP donor template either in ssDNA or dsDNA form, we found that enGagers stimulate significant KI efficiency only for ssDNA but not dsDNA donor. Specifically, the FECO, RecA, GS-FECO and Brex-FECO mRNA enGagers are able to enhance GFP KI to 1.68- to 1.99-fold of a WT Cas9 mRNA 5 days post-electroporation (Fig. 4D-4G). Consistently, when we tested the liposome mediated co-transfection of enGager mRNA/sgRNA together with ssDNA or dsDNA donor templates in HK293 cells, only ssDNA but not dsDNA donors demonstrated 1.35- to 1.99-fold KI enhancement by FECO, Brex-FECO and GS-FECO mRNA enGagers 5 days post-delivery (Fig. 4H&4I). As a reference, WT Cas9 and Cas9-Brex fusion were used as controls in these experiments.

**Fig.4.**
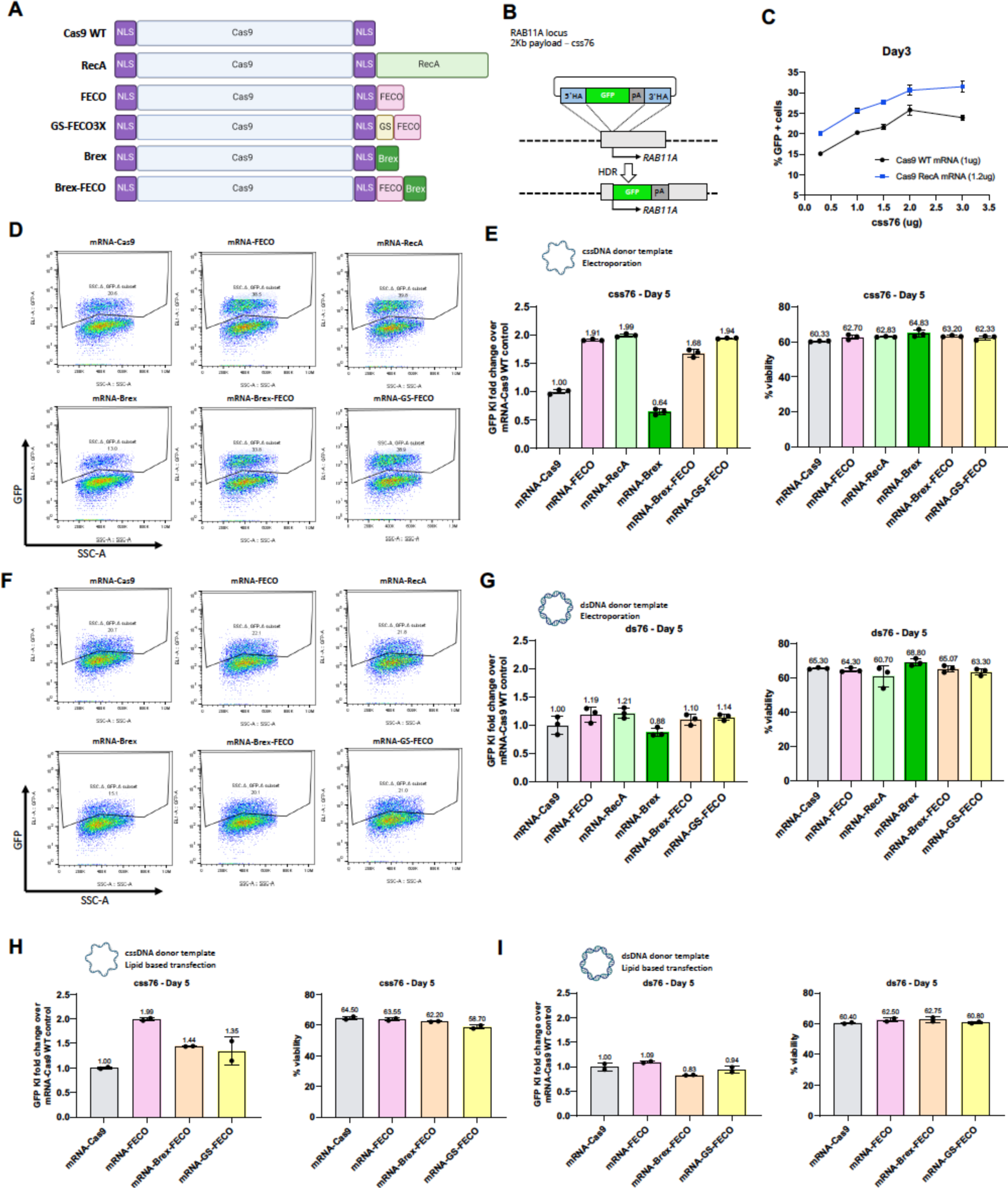
TESOGENASE mediated genome integration enhancement is ssDNA dependent. A, Schematic diagram of various Cas9-ssDNA binding protein and peptide fusion constructs (enGagers) in mRNA form. Two nuclear localization signals were added to the N’ and C’-termini of the Cas9 protein. GS is a shortened multi-GS peptide linker. B, Schematic diagram of Knock in strategy of a 2Kb cssDNA donor template for RAB11A locus. C, Dose titration of cssDNA from 0.3, 1, 1.5, 2 and 3ug for 2Kb GFP transgene knock-in at day 3 post electroporation. Cas9-RecA mRNA enhance around 25-30% knock in efficiency than Cas9 WT mRNA at all the cssDNA dose tested in k562 cells. D, representative FACS profiles with gating strategy showing % of 2Kb GFP transgene cassette Knock-in on RAB11 locus by cssDNA donor at day 5 post electroporation for various enGagers listed in A in K562 cells. E. Quantification of 2Kb GFP transgene cassette Knock in fold change (left) and cell viability (right) of various enGagers as compared to Cas9 WT at day 5 post electroporation from D. F, representative FACS profiles with gating strategy showing % of 2Kb GFP transgene cassette Knock-in on RAB11 locus by dsDNA donor at day 5 post electroporation for various enGagers listed in A in K562 cells. E. Quantification of 2Kb GFP transgene cassette Knock in fold change (left) and cell viability (right) of various enGagers as compared to Cas9 WT at day 5 post electroporation from G. H, Quantification of 2Kb GFP transgene cassette Knock in fold change (left) and cell viability (right) of various enGagers mRNA as compared to Cas9 WT mRNA with cssDNA donor at 5 days post-delivery in HEK293 cells by lipofectamine 3000 transfection. I, Quantification of 2Kb GFP transgene cassette Knock in fold change (left) and cell viability (right) of various enGagers mRNA as compared to Cas9 WT mRNA with dsDNA donor at 5 days post-delivery in HEK293 cells by lipofectamine 3000 transfection. Bars represents mean ± SD from 3 biological replicates.

### TESOGENASE in mRNA form enhance ultra-large transgene integration on various genomic loci

We then assessed the effect of enGagers more broadly on various genomic loci and for donor payload with various size, especially 4 and 8Kb payloads that are around AAV6 and lentivirus packaging limit, respectively. Utilizing highly purified mRNA form of FECO and RecA enGagers, we tested the efficiency of GFP transgene KI on two genomic loci with various length of payload. For *RAB11A* target locus, we designed 2Kb, 4Kb and 8Kb cssDNA payloads, for another target site, the clinically relevant immune cell therapy *B2M* locus (Ren et al., 2017), we designed 2Kb and 4Kb cssDNA payloads (Fig.5A&5D). As illustrated in Fig. 5B, when mRNA enGager/sgRNA with cssDNA donor templates were co-electroporated to K562 cells, FECO and RecA enGagers increase the GFP transgene KI to 44.6-48.5% from 30.10% by WT Cas9 for 2Kb *RAB11A* GFP payload, consistent from previous results. For 4Kb payload, FECO and RecA enGagers increase the GFP transgene KI to 11.07-13.77% from 6. 17% by WT Cas9. For 8Kb payload, FECO and RecA enGagers increase the GFP transgene KI to 3.73-5.17% from 2.97% by WT Cas9. Similarly, for *B2M* locus, FECO and RecA enGagers increase the KI efficiency to 39.6-43.97% from 30.13% by WT Cas9 for 2Kb GFP transgene, and increase the KI efficiency to 10.67-14.07% from 6.27% by WT Cas9 for 4Kb GFP transgene (Fig.5E). As a negative control, Cas9-Brex fusion did not enhance cssDNA mediated transgene KI. Cell viability was not affected by 1 ug Cas9 or enGager mRNA (Fig.5B-C and 5E-F) as compared to Mock electroporation.

**Fig.5.**
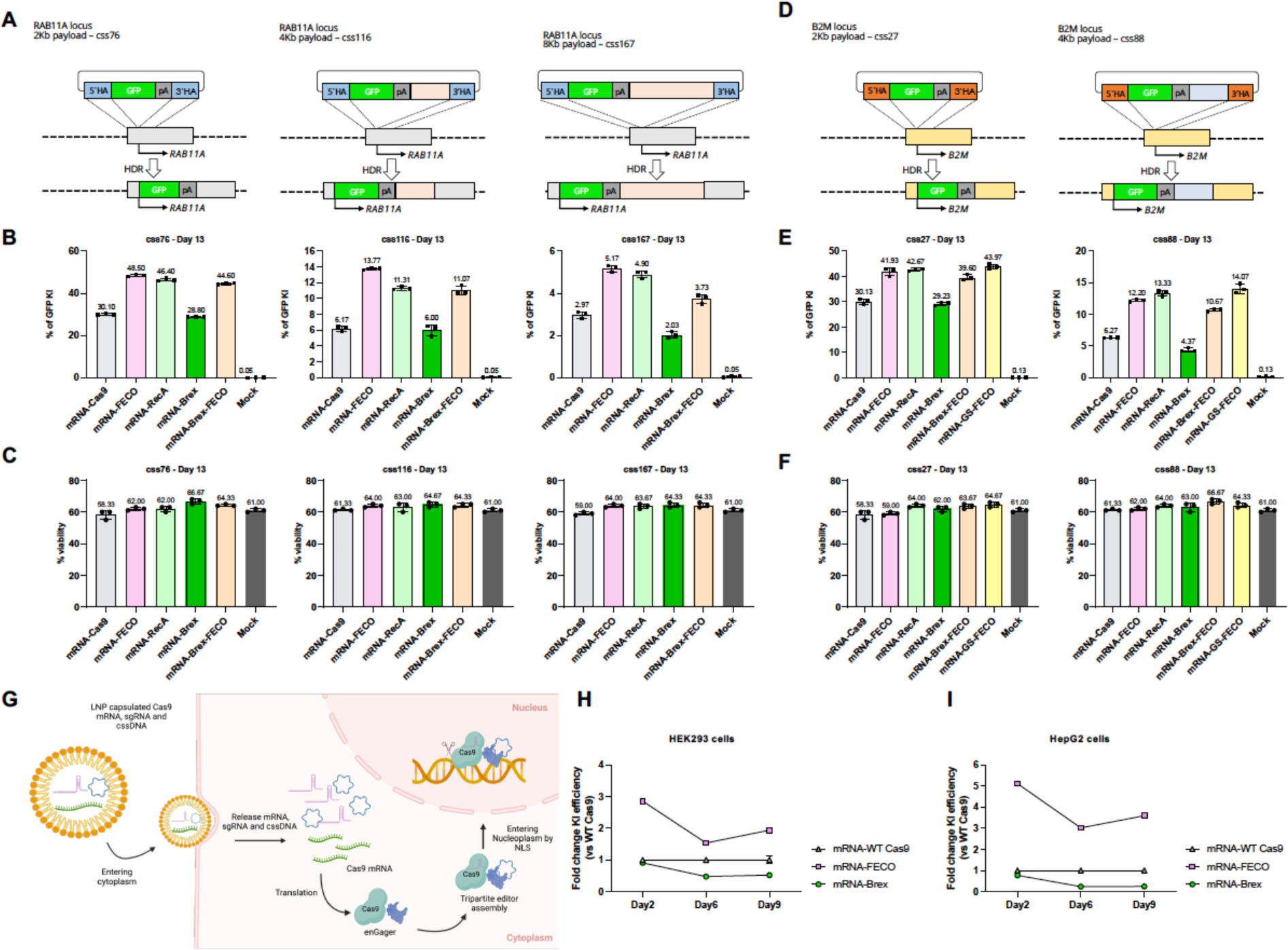
TESOGENASE in mRNA form enhance genome integration on various locus and large payload. A, Schematic diagram of Knock in strategy of 2Kb (css76), 4Kb (css116) and 8Kb (css167) cssDNA donor templates for RAB11A locus. B, Quantification of % of 2Kb (left), 4Kb (middle) and 8Kb (right) GFP transgene cassette Knock-in for various mRNA enGagers RAB11A day 13 post electroporation in K562 cells. C, Quantification of cell viability for 2Kb (left), 4Kb (middle) and 8Kb (right) GFP transgene cassette Knock-in for various mRNA enGagers on RAB11A locus day 13 post electroporation in K562 cells. D. Schematic diagram of Knock in strategy of 2Kb (css27), 4Kb (css88) cssDNA donor templates for B2M locus. E, Quantification of % of 2Kb (left) and 4Kb (right) GFP transgene cassette Knock-in for various mRNA enGagers on B2M locus day 13 post electroporation in K562 cells. F, Quantification of cell viability for 2Kb (left) and 4Kb (right) GFP transgene cassette Knock-in for various mRNA enGagers on B2M locus day 13 post electroporation in K562 cells. Bars represents mean ± SD from 3 biological replicates. G, Schematic diagram showing enGager mRNA/sgRNA/cssDNA delivery into cells using LNP formulation. Once the editor components were delivered into the cytoplasm, the enGager mRNA is translated into endonuclease protein which forms a complex with sgRNA and cssDNA donor template. The assembled tripartite editing machinery complex then can be effectively shuttled into the nucleoplasm and tether onto the target genomic locus for transgene integration. H & I, Quantification of GFP transgene knock-in on RAB11A locus in HEK293 cells (H) and HepG2 cells (I) by LNP delivery at day 2, day 6 and day 9 post-delivery. Data were compared to mRNA-WT Cas9. Bars represents mean ± SD from 2 biological replicates.

### LNP delivery of TESOGENASE and ssDNA donor templates enhance KI

In vivo delivery of Cas9 family endonuclease editing machinery by lipid nanoparticle (LNP) has been successfully demonstrated for gene deletion, base editing and HDR based short gene insertion in rodent and non-human primates (Farbiak et al., 2021; Hou et al., 2021; Musunuru et al., 2021; Qiu et al., 2021; Zhang et al., 2020). Importantly, Gillmore et al has run a clinical trial with an in vivo gene-editing approach to reduce the serum level of toxic TTR as a therapeutic approach to treat Transthyretin amyloidosis (ATTR). This approach is based on a specialized LNP formulation encapsulating mRNA form for Cas9 endonuclease and a sgRNA targeting the disease causing gene *TTR (Gillmore et al., 2021)*. Here we sought to evaluate if the FECO enGager and cssDNA donor template can be co-delivered with LNP potentiating the KI efficiency in various cell types. The hypothesis is that the enGager mRNA/sgRNA/cssDNA can be co-delivered into cytoplasm using LNP formulation. Once in the cytoplasm cytoplasm, the enGager mRNA is translated into endonuclease protein which complexes with sgRNA and cssDNA donor template using its ssDNA binding motif. The assembled tripartite editing machinery complex then can be effectively shuttled into the nucleoplasm and tether onto the target genomic locus for transgene integration (Fig.5G). As shown in Fig.5H&I, when delivered in HEK293T and HepG2 cells with an off-the-shelf liver cell targeting LNP, FECO enGager demonstrated a persistent enhancement of a 2Kb GFP transgene KI on RAB11A locus by 1.5- to 3-fold and 3- to 5-fold from Day 2 to Day 9 post-delivery, respectively. Consistent with previous observation with other delivery approaches, the Brex mRNA reduced the KI efficiency overtime. These data demonstrated that the enGagers can be delivered in mRNA form by ***NanoAssemblr*** LNP for potentiated genome integration and potentially be applied for in vivo genome integration.

### CAR-T engineering by TESOGENASE with superior efficiency than WT SpCas9

Finally, we sought to use the newly developed FECO enGager to engineer primary T cells by inserting a functional chimeric antigen receptor (CAR) transgene cassette on T cell receptor alpha constant chain (*TRAC*) locus (Eyquem et al., 2017). We chose a ∼ 3Kb CD19-CD22 dual CAR construct that has been demonstrated anti-tumor function for potential treatment of patient with Acute Lymphoblastic Leukemia (ALL) (Fig. 6A and Fry et al., 2018). We used PE conjugated Protein-L binder to analyze the KI efficiency. When delivered by co-electroporation with CD19-CD22 dual CAR cssDNA doner template in CD4+/CD8+ primary pan-T cells, GS-FECO mRNA enGager achieved 30.2% to 33.4% targeted CAR KI at day 7 post-delivery, whereas WT Cas9 mRNA only achieved 5% to 14.1% of CAR KI with various mRNA dosages. Especially when lower dose of mRNA editors was used, GS-FECO enGager enhance cssDNA mediated CAR-T engineered by >6 fold (Fig.6B). Importantly, GS-FECO enGager mediated enhancement of dual CAR-T engineering did not compromise T cell counts, proliferation and cell viability compared to WT Cas9 (Fig.6C). To test the anti-tumor function of the engineered CAR-T cells, we use the Incucyte live imaging approach to monitor the cell killing functions over the course of 94 hours. We titrated the E:T ratio at 2:25:1, 4.5:1 and 9:1. When co-cultured with a leukemia lymphocyte NALM6 cells, GS-FECO enGager mRNA engineered CD19-CD22 dual CAR-T cells demonstrated more effective and durable cancer cell killing function compared to WT Cas9 mRNA engineered dual CAR-T cells at every E:T ratio we tested (Fig.6D&6E)

**Figure 6.**
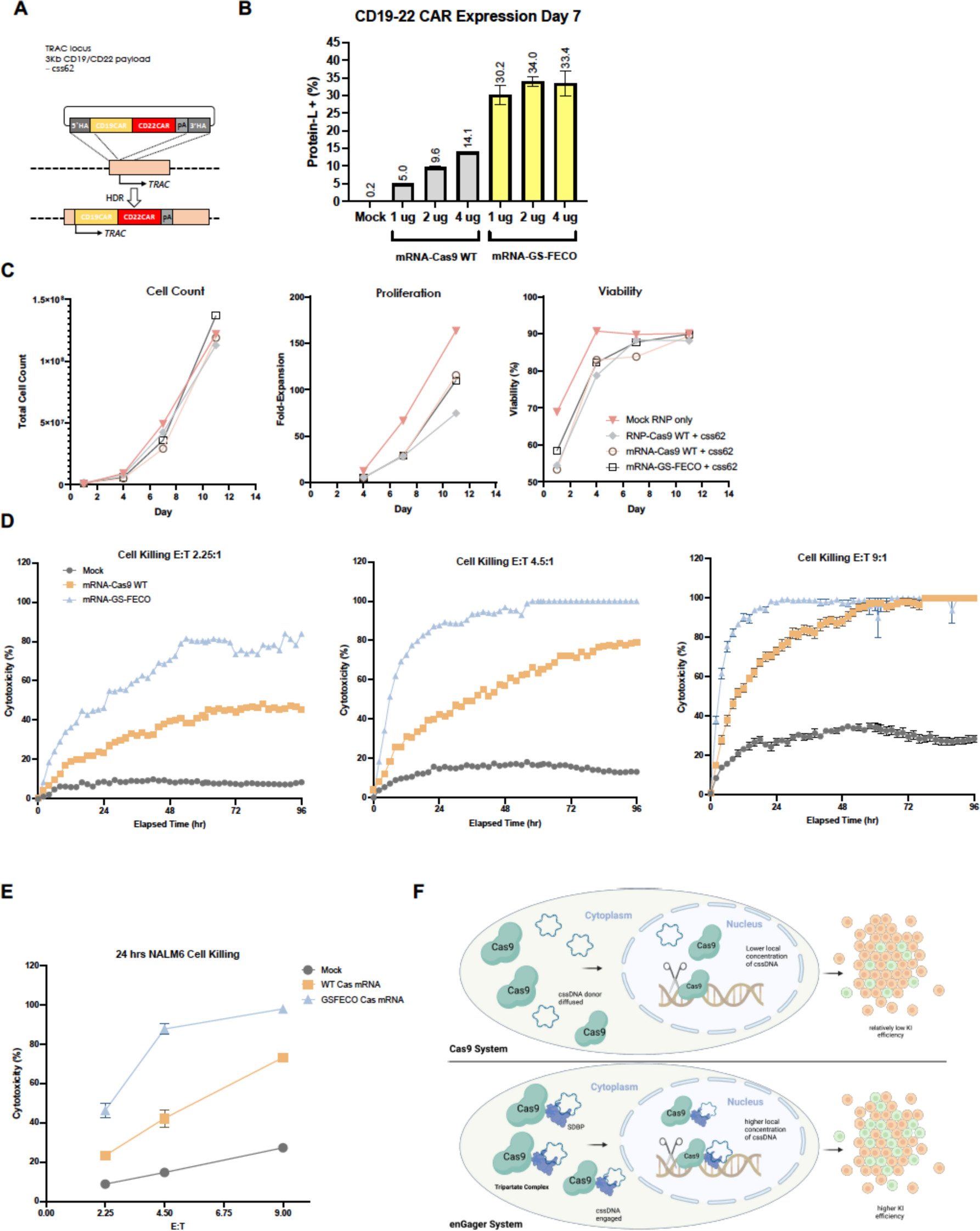
CAR-T engineering by enGager with superior efficiency than WT Cas9. A, Schematic diagram of Knock in strategy of 3Kb CD19/CD22 dual CAR (css62) cssDNA donor templates for TRAC locus in primary T cells. B, Quantification of % of CD19/CD22 CAR Knock-in using cssDNA donor analyzed by CD19 binder for various doses of Cas9 WT and Cas9-GS-FECO enGager mRNA at 1 ug, 2ug and 4ug. Data were collected at day 7 and day 11 post electroporation of primary T cells. Cas9-GS-FECO enGager achieves ∼ 4- to 6-fold higher CAR-T engineering efficiency than Cas9 WT. C, Characterization of the engineered CAR-T cells or mock treated T cells for total cell count, cell proliferation fold change and cell viability over time. Bars represents mean ± SD from 2 biological replicates. D, NALM6 leukemia lymphocyte killing curve of unengineered T cells, CD19-CD22 dual CAR-T cells engineered with 2 ug of WT Cas9 mRNA and 2 ug of GS-FECO enGager mRNA over the course of 96 hrs. Effect (T cells): Target (NALM6 cells) are at 2.25:1 for left panel, 4.5:1 for middle panel and 9:1 for right panel. E, NALM6 cell killing function of CAR-T cells at 24 hrs for E:T ratio at 2.25:1, 4.5:1 and 9:1. F, Schematic diagram of engineered enGagers with single stranded DNA binding protein (SSBP) can recruit cssDNA donor template and form a tripartite editing machinery for efficient translocation of the entire editing complex from cytoplasm to nucleus. As a result, the donor DNA has higher effective local concentration in the nucleus for more efficient homologous directed genome integration. This process works more prominently with cssDNA.

## DISCUSSION

In this study, we presented a set of enGager endonucleases by fusing Cas9 with various full-length single-stranded DNA binding protein modules or their corresponding miniature motifs with up to 20 amino acids. When applied for targeted genome integration with cssDNA donor templates to diverse genomic loci across multiple cell types, these enGagers outperform conventional editors by up to several times for small and large transgene knock-in. Using one of the small enGagers, we demonstrated sizeable chimeric antigen receptor (CAR) transgene integration in primary T cells with exceptional efficiency and anti-tumor cell function. As shown in Fig.6F schematic diagram, the engineered enGagers with single-stranded DNA binding protein (SSBP) can recruit cssDNA donor template and form a tripartite editing machinery for efficient translocation of the entire tripartite editing complex from cytoplasm to the nucleoplasm. As a result, the donor DNA has a higher effective local concentration in the nucleus and to the targeted genomic locus for more efficient homologous directed genome integration (bottom). In contrast, Cas9 alone cannot engage and recruit cssDNA donor templates; therefore, the donor DNA templates are restricted in the cytoplasm with a relatively low level of gene KI efficiency. Additionally, enGagers increase KI efficiency for single-stranded DNA template but not double-stranded DNA because SSBP fused to enGager is unable to recruit double-stranded DNA. This difference between single-stranded DNA and double-stranded DNA further validates tripartite complex mechanism. These newly designed enGager editors expanded the gene-editing toolbox for potential cell and gene therapeutic development based on ssDNA-mediated non-viral genome engineering.

Previously, it was challenging to achieve gene knock-in for >4kb payload especially for non-viral knock-in approaches. This makes gene knock-in improvement by enGager more attractive for many therapeutical applications that require large inserts to be functional. Additionally, the efficacy of enGagers was tested against different variables including multiple cell types, loci and payload sizes, and enGagers exhibited consistent improvement in knock-in efficiency. This improvement also includes an increase in efficiency for extremely large 8kb payload from ∼3% to ∼5%. Since many previous studies struggled to achieve meaningful knock-in for >4 kb payload, our finding may open new opportunities for Cas9 mediated precise gene therapy field. These new opportunities include multiple or large gene knock-in and long regulatory elements knock-in along with target gene. enGager also improve knock-in of smaller 2 kb payload, therefore the benefit of the enGager is not just limited to large payload. Also, unlike AAV viral mediated delivery, enGager delivers HDR payload to nucleus without the concerns of viral toxicity and unwanted integration while being relatively easy to manufacture.

One of the most critical findings of this study was that the 20aa single-stranded DNA binding motif of RecA showed comparable improvement relative to that of the full-length RecA protein. This observation underscores our hypothesis that the formation of tripartite complex rather than the recombination activity of RecA contributes to knock-in efficiency. We reason that the enGager platform can benefit from the 20aa motif over full-length proteins in several ways. First, the 20aa motif fusion enGagers are smaller than full protein fusion enGagers. spCas9 protein is a relatively large protein, with a molecular weight of approximately 160kDa. Further increasing the size of Cas9 may hamper encapsulation, delivery and manufacturing. In contrast, the 20aa motif is very small in size and less likely to produce such problems. Second, since the 20aa motif only has single-stranded DNA binding capacity, it is less likely to have unwanted side effect than full size protein that has more functionality. Third, the 20aa motif fusion is extremely small and could be further improved by adding additional fusion protein that has different mechanisms of action.

While enGagers demonstrated significant improvement in gene knock-in efficiency, there are still some limitations of enGagers that calls for further studies. While 4kb payload showed meaningful knock-in efficiency, 8kb knock-in efficiency only showed up to 5% knock-in. Although relative improvement of enGagers from Cas9 only construct is similar to 2kb and 4kb payload, absolute knock-in rate for 8kb payload is still too low to be meaningful. Therefore, further engineering is required to claim effective 8kb payload knock-in. Additionally, our study doesn’t include in-vivo study because we mostly utilized electroporation transfection method which is limited to ex-vivo or in-vitro usages. Lipofection was also tested and showed promising result but for more robust delivery platform such as lipid nanoparticles (LNP) can further improve enGager platform especially for potential in-vivo delivery. The follow-up study will aim to address these limitation to further improve the enGagers platform. For 8kb payload, we aim to further test different fusion motif candidates that might show higher improvement. Also, we plan to fuse additional protein that improve HDR through different mechanism of action. For transfection, we seek to test our mRNA platform with LNP. This future study could include potential in-vivo delivery of enGager mRNA and circular ssDNA HDR template.

In summary, enGager was able to significantly improve the KI efficiency of various donor templates up to several folds in various cell types. One of the most advantageous aspects of enGager was its size being smaller than 20aa long, allowing enGager to be compact and opens possibility of further improvement by additional components. Also, enGager improved 8kbp KI but still was unable to increase the efficiency enough for practical applications. We also need to expand and test different transfection platform such as LNP. Despite these limitations, enGager and its compact fusion platform will likely improve both KI platform development and practical applications like CAR-T cell engineering. We hope to also see enGager can be further improved to raise 8kbp KI to practical level while expand to in-vivo transfection platform like LNP delivery in the future.

## METHODS and MATERIALS

### Plasmid constructs

For all-in-one (AIO) plasmid construct design for various enGagers (Table 1), ssDNA binding protein and peptide sequences were synthesized and assembled into an AIO plasmid modified from Addgene plasmid #42230 by golden gate cloning. Complex fusion constructs were cloned using multi-piece DNA Gibson assembly approach. For mRNA constructs, various enGager coding sequences were cloned into a modified vector plasmid from pGEM®-4Z vector containing T7 promoter, 5’ UTR, Cas9 fusion sequence ORF 3’ UTR, bovine growth hormone polyadenylation signal (bGH polyA), and 64 poly adenine sequence (Promega). Table 1 listed the amino acid sequences of the enGager fusion protein and WT Cas9 endonuclease.

### mRNA production and purification

mRNA in this study was produced through in vitro transcription from plasmid templates. Frist plasmids containing T7 promoter, 5’ UTR, Cas9 fusion sequence ORF 3’ UTR, bovine growth hormone polyadenylation signal (bGH polyA), and 64 poly adenine sequence were cloned. To linearize the DNA, Spe1 restriction site was inserted at the end of the 64 poly adenines. After plasmids were produced, ∼50µg of plasmids were digested using a restriction enzyme to cut the plasmids at the added restriction site for linearization. After 24 hours, the resulting linear DNA was cleaned through a DNA clean-up kit from zymo (Cat# D4029). After DNA cleanup, the linear DNA fragments were introduced to HiScribe™ T7 mRNA Kit with CleanCap® Reagent AG (Cat# E2080S) from New England Biolabs. This kit used AG CleanCap® Reagent from TriLink Biotechnologies and yielded ∼ 100µg per 1µg of linear DNA templates. For each reaction, 1µg of templates were mixed with 2µl 10x reaction buffer, 2µl of ATP (60mM), 2µl of GTP (50mM), 2µl of UTP (50mM), 2µl of CTP (50mM), 2µl of Cap Analog (40mM) and 2µl of RNA polymerase Mix. Then Nuclease-free water was added to the final volume of 20µl to the mixtures. The mixture was then incubated at 37C° for 2 hours. After 2 hours of in vitro transcription, the resulting mRNA was purified using RNA Clean & concentrator from Zymo (Cat # R1019). DNase was also used during the purification to remove residual DNA templates from the solutions. After in vitro transcriptions, each mRNA was analyzed using agarose gel analysis. 500ng of mRNA were diluted to 20ul of nuclease-free water then 40ul of 2 x RNA loading dye from ThermoFisher Scientific (Cat# R0641) was added. The mixture was then incubated at 80C° for 15 minutes. After incubation, the mRNA mixtures were immediately transferred to Ice and ran on the clean 1% agarose gel made from TAE buffer.

### LNP formulation and Delivery

1. HEK293T and HepG2 cells were plated on 24 well format day, a day prior to transfection in 500uls of antibiotics free and reduced serum (5%) DMEM media. 150k cells were plated / well of a 24 well plate
2. On the day of electroporation, cells were pretreated with 5uls of APoE (100X)
3. Using NanoAssemblr® Spark™ system (Precision NanoSystems), 7 different formulations were prepared following the manufacturer protocol:

- Formulation 1# Cas9 mRNA + RAB11A sgRNA + cssDNA
- Formulation 2# FECO mRNA + RAB11A sgRNA + cssDNA
- Formulation 3# Brex mRNA + RAB11A sgRNA + cssDNA
4. After preparation, formulations were diluted at 1:1 ratio.
5. LNP formulations were administered in 1:1 ratio in combination with css76 formulations
6. Cells were dissociated for FLO assays at day 2, day 6 and day 9 post-delivery.

### HEK293 lipofectamine transfection

For lipofectamine transfection of mRNA editors/sgRNA (mRNA cocktail) and DNA donors, 24 well plates are coated with PLO for 2hrs. Plates are washed 2X and dried before they were plated with cells. For each well 2×10^5^ cells were plated on Day 0 using 500 ul of complete DMEM media. On Day 1, 250uls of media was slowly replaced with equal volume of serum free and antibiotic free DMEM media. Both mRNA cocktail and DNA were prepared separately as per the conditions. To prepare the mRNA cocktail, individual mRNA construct (1ug/well) and Rab11A sgRNA (2.54ul/well from a stock of 80uM) were diluted in 1:1 ratio with lipofectamine 3000 and incubated on ice for 10-15 minutes; for DNA prep, both single stranded and double stranded DNAs were packaged separately. For both, concentration of 1ug/well was used. Respective DNA was mixed with P3000 reagent (1ug/well) and the whole mix was diluted with lipofectamine 3000 in 1:1 dilution. The DNA cocktail was also incubated at RT for 10-15 mins. After 15 mins of incubation, mRNA cocktail and DNA cocktail were added in 1:1 ratio as per the plate map. Cells were then allowed to sit for 48 hours after which they were transferred from 24 well plate format to 6 well plate with 1ml media. FLO analysis was performed on D5, D9 and D14 to check on the knock in efficiency of the constructs.

### Generation of circular single-stranded DNA

Donor template sequences (transgene sequence flanked with 5’ and 3’ homology arms at 300-500 nt in length) are constructed as dsDNA and placed into phagemid vector by Golden Gate Assembly. An XL1-Blue E. Coli Strain was co-transformed with the M13 helper plasmid and phagemid containing donor template sequences and selected in agar plate with kanamycin (50 µg/mL), carbenicillin (100 µg/mL). Single colony was selected and grown for ∼24 hours (37 °C, 225 rpm) in 250 mL 2xYT media (1.6% tryptone, 1% yeast extract, 0.25% NaCl) to reach OD600 between 2.5-3.0. The bacteria were pelleted by centrifugation and the phage particles in the supernatant were precipitated by PEG-8000 by centrifugation, washed and lysed in 20 mM MOPS., 1M Guanidine-HCl and 2% Triton X-100. The cssDNA released from phage were then extracted with NucleoBond Xtra Midi EF kit (Macherey-Nagel) following the manufacturer’s instructions. The concentration of cssDNA is determined by Nanodrop. Ratios of Absorbance (A260 nm/280 nm and 260nm/230nm) will reflect consistent purity (1.8 and > 2, respectively) from serial preps. Recombinant cssDNA is verified by DNA sequencing using custom-designed staggered sequencing primers for complete coverage.

### Cell culture

K562 cells (ATCC, CCL-243) were maintained in RPMI-1640 media with 10% FBS and 1% penicillin and streptomycin. HEK293T (ATCC) cells were cultured in Dulbecco’s modified Eagle’s medium supplemented with 10% fetal bovine serum and 1% penicillin and streptomycin (Gibco). HepG2 cells (HB-8065™) were cultured in ATCC-formulated Eagle’s Minimum Essential Medium (Catalog No. 302003) supplemented with 10% FBS. iPSC (Thermo Fisher) was cultured in StemFlex (Thermo Fisher) media in vitronectin-coated flask. iPSC colonies were checked regularly and passaged using ReLeSR (StemCell Technologies) every 3-4 days of culture. iPSCs were ready for electroporation after 2-3 passages. T cells were cultured and expanded in TexMACS Medium (Miltenyi) supplemented with 200 IU/mL Human IL-2 IS (Miltenyi). T cells were activated for 2 days with T Cell TransAct (Miltenyi) before electroporation. All cells were maintained in a humidified incubator at 37 °C and 5 % CO2, unless otherwise specified. Cell count viability were determined using Via2-Cassette in NucleoCounter® NC-202 (ChemoMetec) on specified days after engineering.

### Electroporation of Cas9 ribonucleoprotein, AIO plasmid and mRNA/sgRNA complex with DNA donor

All K562, HEK293T, iPSC and T cell electroporation were performed using the Amaxa™ 96-well Shuttle™ with the 4D Nucleofector (Lonza). Cas9 nucleases and sgRNAs were precomplexed in supplemented Nucleofector® Solution for 20 min at room temperature and the RNP solution was made up to a final volume of 2.5 µL (10X). For electroporating K562 cells, SF Cell Line 4D-Nucleofector™ Kit and 250,000-500,000 cells per reaction were used with program FF-120. For electroporating HEK293T cells, SF Cell Line 4D-Nucleofector™ Kit and 200,000-300,000 cells per reaction were used with program FS-100. For electroporating iPSC cells, 100,000 cells per reaction were used with P3 Primary Cell 4D-Nucleofector™ Kit and program CA-137. For electroporating primary T cells, 2×10^6^ cells per reaction were used with P3 Primary Cell 4D-Nucleofector™ Kit and program EO-115. Indicated amount of HDR donor template (cssDNA or dsDNA) were co-electroporation with RNP. For mRNA or AIO plasmid enGager electroporation, 1 ug-1.5ug of mRNA or DNA were electroporated together with indicated amount of HDR donor templates, with the same electroporation parameters used for RNP electroporation. After electroporation, cells were incubated in humidified 32°C incubator with 5 % CO2 for 12-24 hours followed up transferring to 37°C incubator for additional days.

### Flow Cytometry analysis

All flow cytometry was performed on an Attune NxT flow cytometer with a 96-well autosampler (ThermoFisher Scientific). Unless otherwise indicated, cells were collected 3-14 days post electroporation, resuspended in fluorescence-activated cell sorting (FACS) buffer (2% BSA in PBS) and stained with 7-AAD (BioLegend) as a dye for cell viability assay, and the indicated cell-surface marker, such as PE labelled Protein L (ACROBiosystems), or FITC-Labeled Human CD19 (20-291) Protein, Fc Tag (ACROBiosystems). To obtain comparable live cell counts between conditions, events were recorded from an equivalent fixed volume for all samples. Data analysis was performed using FlowJo_v10.8.0_CL software with exclusion of subcellular debris, singlet gating and live/dead staining. Data analysis is performed using FlowJo_v10.8.0_CL software. Data were plotted using Prism Graphpad.

### CAR-T cell engineering

Primary T cells were isolated and enriched from Leukopak using Pan T Cell MicroBead Cocktail with MultiMCAS Cell24 Separator Plus (Miltenyi), CD4+ and CD8+ pan T cells were cryopreserved for later use. T cells were cultured and expanded in TexMACS Medium (Miltenyi) supplemented with 200 IU/mL Human IL-2 IS (Miltenyi). T cells were activated for 2 days with T Cell TransAct (Miltenyi) before electroporation. 48 h after initiating T-cell initiation and activation, T cells were electroporated using Amaxa™ 96-well Shuttle™ in 4D Nucleofector. 2×10^6^ cells were mixed with 25 pmol of Cas9 WT protein or 1 ug of enGager mRNA and 50 pmol of gRNA into each well. For cssDNA engineered cells, 2 µg of cssDNA encoding bi-specific CD19×CD22 CAR (Fry et al., 2018) was electroporated with RNP or mRNA/sgRNA cocktail targeting TRAC locus. Following electroporation, cells were diluted into culture medium in the presence of T Cell TransAct with 1 µM DNA-PK inhibitor M-3814, and incubated at 32 °C, 5% CO2 for 24 hours. Cells were then washed and subsequently transferred into G-Rex 24 Multi-Well Cell Culture Plate (Wilson Wolf) in standard culture conditions at 37 °C, 5% CO2 in IL-2 supplemented TexMACS medium and replenished every 3-4 days. CAR expressions were determined using PE labelled Protein L (ACROBiosystems), or FITC-Labeled Human CD19 (20-291) Protein, Fc Tag (ACROBiosystems).

## Author Contributions

H.W. and R.S. conceived the idea for this project. H.N., K.X. and H.W. designed the experiments and interpreted the data. H.N., K.X., I.M., S. B, J.S. and D.L. performed the experiments. J.L. and H.W oversaw the study. H.N. and H.W. wrote the manuscript with inputs from all the other authors. I.M., J.S., D.L., K.X., R.S. and H.W. are paid employee from Full Circles Therapeutics, Inc.

## ACKNOWLEDGMENTS

We thank Quintara Bioscience Inc. for plasmid construction, and DNA/RNA sequencing service for the study. J.L. receives funding from the Northeastern University start-up, National Institute of Biomedical Imaging and Bioengineering (R21EB030769), National Science Foundation CAREER (2238972), National Institute of Dental and Craniofacial Research awards (R03DE031329 and R01DE030943), CDMRP PRCRP Idea Award (W81XWH-21-1-0324) and CDMRP PRMRP Discovery Award (W81XWH-21-1-0016).

